# Heat-off responses of epidermal cells sensitize *Drosophila* larvae to noxious inputs

**DOI:** 10.1101/2025.05.30.656957

**Authors:** Jiro Yoshino, Avery Chiu, Takeshi Morita, Chang Yin, Federico M. Tenedini, Takaaki Sokabe, Kazuo Emoto, Jay Z. Parrish

**Affiliations:** Department of Biology, University of Washington, Campus Box 351800, Seattle, WA 98195, USA; Department of Biological Sciences, Graduate School of Science, The University of Tokyo, 7-3-1 Hongo, Bunkyo-ku, Tokyo 113-0033 Japan; Laboratory of Neurogenetics and Behavior, The Rockefeller University, 1230 York Avenue, New York, NY 10065; Thermal Biology Group, Exploratory Research Center on Life and Living Systems, National Institutes of Natural Sciences, Okazaki, Japan; Section of Sensory Physiology, Center for Genetic Analysis of Behavior, National Institute for Physiological Sciences, National Institutes of Natural Sciences, Okazaki, Japan; Graduate Institute for Advanced Studies, SOKENDAI, Hayama, Japan.; International Research Center for Neurointelligence (WPI-IRCN), The University of Tokyo, 7-3-1 Hongo, Bunkyo-ku, Tokyo 113-0033 Japan

**Keywords:** epidermis, somatosensory neuron, nociception, thermosensation, sensitization

## Abstract

Perception of external thermal stimuli is critical to animal survival, and although an animal’s skin is the largest contact surface for thermal inputs, contributions of skin cells to noxious temperature sensing have not been extensively explored. Here, we show that exposure to heat transiently sensitizes *Drosophila* larvae to subsequent noxious stimuli. This sensitization is induced by prior stimulation of epidermal cells but not nociceptors, suggesting that epidermal cells modulate nociceptor function in response to heat exposure. Indeed, we found that *Drosophila* epidermal cells are intrinsically thermosensitive, exhibiting robust heat-off responses following warming to noxious temperatures as well as responses to cooling below comfortable temperatures. Further, we found that epidermal heat-off calcium responses involve influx of extracellular calcium and require the store-operated calcium channel Orai and its activator Stim. Finally, epidermal heat-off responses and heat-evoked nociceptive sensitization exhibit similar temperature dependencies, and we found that Stim and Orai are required in epidermal cells for heat-evoked nociceptive sensitization. Hence, epidermal thermosensory responses provide a form of adaptive sensitization to facilitate noxious heat avoidance.

## Introduction

Animals encounter a broad range of temperatures in their environment and therefore have evolved robust mechanisms to sense temperature changes, discriminate between comfortable and noxious temperatures, and avoid the latter. Temperature-guided behaviors are particularly important for ectotherms, which rely on these behaviors for thermoregulation and for limiting exposure to temperature extremes that adversely affect performance and fitness^1^. For example, extended exposure to stressful heat or cold significantly reduces viability, fecundity, progeny lifespan, and increases variability in a variety of quantitative traits in many *Drosophilidae*^2,3^.

*Drosophila* larvae exhibit thermally guided behaviors that are mediated by multiple temperature sensing pathways, with distinct subsets of larval sensory neurons mediating escape from noxious temperatures and thermotaxis towards favorable temperatures^4–7^. Exposure to temperatures > 40°C evokes nocifensive corkscrew-like rolling behavior in larvae^7^, and inactivating class IV dendrite arborization (C4da) neurons strongly attenuates this response, suggesting that C4da neurons are the primary noxious heat sensors. Indeed, C4da neurons directly detect noxious heat via expression of TRP channels including Painless, TRPA1, and Pyrexia^7–9^, and optogenetic stimulation of C4da neurons triggers nocifensive rolling behavior in larvae^8^. However, the larval responses to noxious heat are subject to modulation by central and peripheral signals. Centrally, descending inputs modulate the activity of nociceptive circuits, as in vertebrates^10^, and starvation triggers release of insulin-like peptides that activate GABAergic inputs to nociceptors^11^. Likewise, sensory inputs during development modulate nociceptor output via serotonergic signaling^12^. In addition to central modulation, peripheral signaling pathways contribute to nociceptive sensitization. For example, larval exposure to UV irradiation triggers thermal allodynia and hyperalgesia via the combined activity of several pathways including activation of Hedgehog and Tachykinin signaling in nociceptors and inflammatory cytokine signaling from epidermal cells^13–15^. Systemic exposure to vinca alkaloids triggers both thermal hyperalgesia and mechanical allodynia by TRPA1-dependent hyperexcitability of C4da neurons^16^.

Although much is known about the neural substrates for noxious temperature sensing, roles for skin cells in noxious heat sensing and avoidance responses have not been extensively studied, even though an animal’s skin is its largest contact site for environmental stimuli. Skin cells have historically been thought of as passive participants in somatosensation, indirectly influencing sensory transduction as filters for sensory stimuli^17^, via effects on somatosensory neuron (SSN) morphogenesis^18^, as accessory cells in specialized end organs^19^, or by the release of inflammatory mediators following tissue injury or disease^20^. However, targeted stimulation of skin cells elicits SSN activity and a range of animal behaviors, including nociceptive escape responses in mice and flies^21–24^. Furthermore, a growing body of evidence demonstrates that resident skin cells actively sample environmental stimuli including temperature. For example, mouse keratinocytes express thermosensitive ion channels^25–28^, are intrinsically responsive to cooling^29,30^ and the calcium release activated calcium (CRAC) channel Orai and its activator Stim act in epidermal cells to regulate thermal preference in mice^30^. While these findings highlight the capacity for epidermal cells to sense external temperature, their unique contribution to thermosensation — distinct from that of somatosensory neurons — remains poorly understood. Moreover, whether epidermal cells contribute specifically to nociceptive temperature detection has yet to be explored.

Here, we examined the capacity of *Drosophila* epidermal cells to sense changes in environmental temperature and modulate animal behavior accordingly. Epidermal mechanosensory responses trigger a form of short-term mechanically-induced nociceptive sensitization^23^, and we found that *Drosophila* larvae likewise exhibit acute sensitization to noxious heat that is uniquely mediated by epidermal stimulation. Our *in vitro* and *ex vivo* calcium imaging experiments demonstrate that epidermal cells are intrinsically thermosensitive, exhibiting robust heat-off responses but minimal responses to warming stimuli. Finally, we find that the epidermal heat-off responses and heat-evoked nociceptive sensitization require the store-operated calcium channel Orai, and its activator Stim.

## Results

### Epidermal cells sensitize larvae to noxious heat

*Drosophila* larvae presented with two successive nociceptive mechanical stimuli exhibit enhanced behavioral responses to the second stimulus^31^ as a result of mechanically-evoked epidermal responses that yield prolonged sensitization to noxious mechanical inputs^23^. Noxious heat triggers nocifensive rolling responses^7^, and thermal hyperalgesia induced by UV damage is manifest as reduced rolling latency^13^, therefore we examined whether a prior heat stimulus impacted rolling latency when larvae were presented with noxious heat. For these assays we fixed the noxious heat stimulus at 41.5°C, a temperature that yielded nocifensive rolling responses in >75% of larvae within 8-10 s of the first stimulus^32^ (Fig. 1A). To control for impacts of handling involved with delivery of the thermal stimulus, we assayed for effects of prior exposure to a 25°C stimulus on rolling latency. Compared to controls that received no prior stimulus, prior exposure to a 25°C stimulus had no significant effect on the probability or latency of heat-induced rolling responses (Fig. 1B, 1C). In contrast, we observed significant decreases in rolling latency when larvae had prior exposure to thermal stimuli ranging from 30°C to 45°C. These effects were graded with temperature: within this thermal range, increasing temperatures of the prior stimulus yielded increased response rates and reduced latencies. Prior exposure to extreme heat (45°C) induced transient paralysis in some larvae, accounting for the reduced rolling probability, and had intermediate effects on rolling latency in responding larvae. Hence, exposure to prior heat sensitizes larvae to noxious thermal inputs, but efficacy of the prior stimulus is limited by an upper threshold of < 45°C.

**Figure 1.**
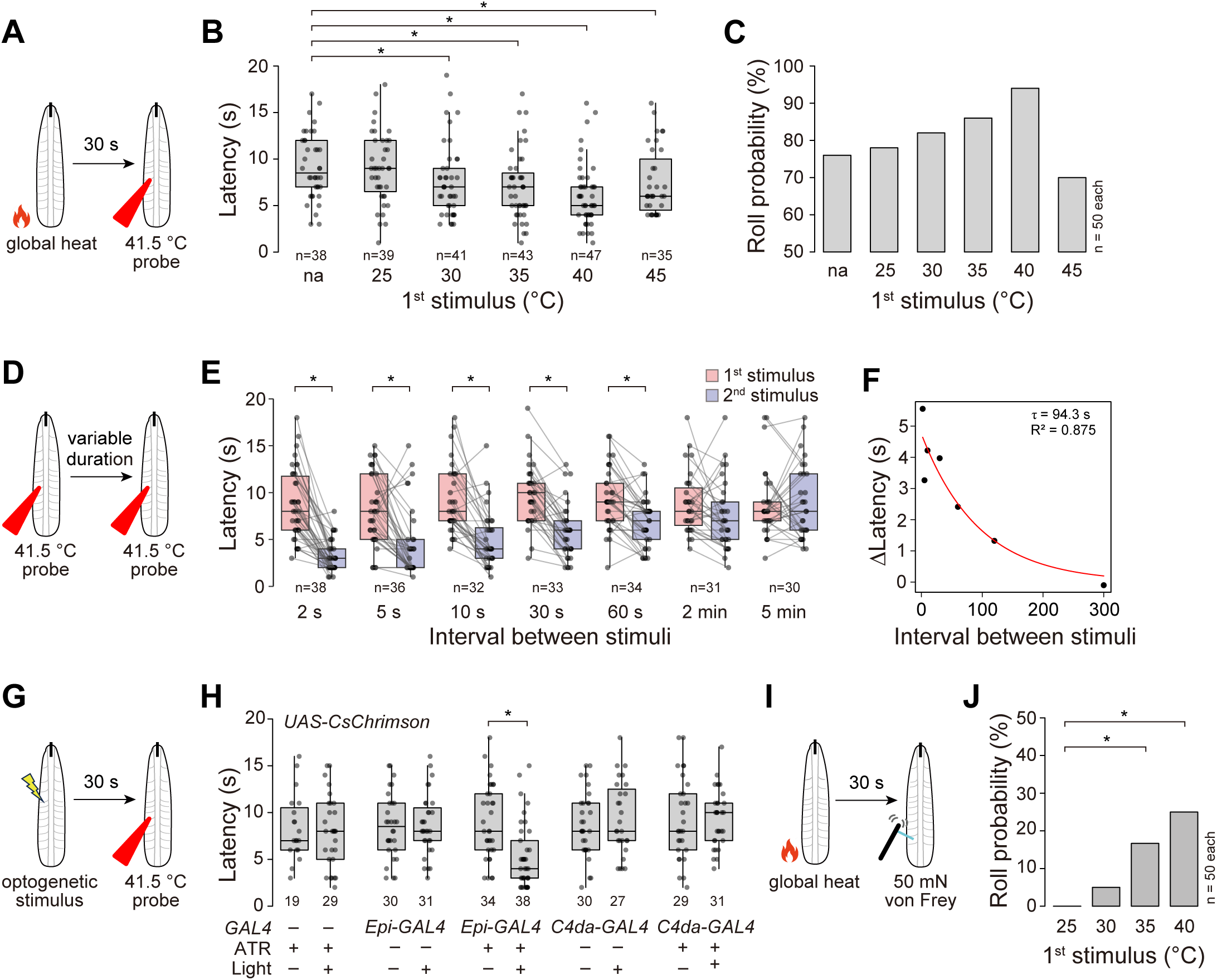
Epidermal stimulation augments thermal nociceptive responses. (A-C). Drosophila larvae exhibit heat-evoked nociceptive sensitization. (A) Schematic of experimental paradigm. Control larvae (w*^1118^*) received a global heat stimulus of varying temperature (25°C – 45°C) on a pre-warmed peltier plate. Following 30 s of recovery they received a 41.5°C stimulus with a thermal probe. (B) Temperature threshold for heat-induced sensitization. Prior thermal stimuli from 30°C to 45°C sensitizes larvae to a subsequent noxious thermal stimulus. *P < 0.05 compared to larvae receiving no prior stimulus; Kruskal-Wallis test followed by Wilcoxon rank sum test with BH correction. (C) Effects of prior exposure to noxious heat on the roll probability to a second stimulus. (D-F) Kinetics of heat-induced nociceptive sensitization. (D) Control larvae (w^11^^18^) received two successive 41.5°C stimuli separated by the indicated recovery period. (E) A 41.5°C stimulus delivered between 2 and 60 s prior to the second stimulus yielded a significant reduction in rolling latency. The plot depicts roll latency to each of the two 41.5°C stimuli spaced by the indicated recovery duration. In this and subsequent box plots, points represent measurements from individual samples, boxes display the first and third quartiles, hatches mark medians, and whiskers mark 1.5 times the interquartile range. *P < 0.05, paired Wilcoxon rank sum test with BH correction. (F) Nociceptive enhancement (difference in the roll latency to the first and second stimulus) is plotted against the recovery duration, and results were fit to an exponential curve f(x) = a*exp(b*x) using R software to derive the decay time constant (τ = 94.3 s). (G) Prior epidermal stimulation sensitizes larvae to noxious heat. Plot depicts roll latency of effector controls (w^1118^; UAS-CsChrimson/+), larvae expressing CsChrimson in the epidermis (w^1118^; R38F11-GAL4, UAS-CsChrimson/+), and larvae expressing CsChrimson in C4da neurons (w^1118^; ppk-GAL4, UAS-CsChrimson/+) in response to 41.5°C thermal stimulus. Larvae were raised on media containing (+ ATR) or lacking (-ATR) the obligate CsChrimson co-factor ATR, as indicated. To control for effects of genetic background, we confirmed that each of the experimental genotypes exhibited heat-induced nociceptive sensitization (Supplementary Fig. 1). *P < 0.05 Wilcoxon rank sum test with BH correction. (I-J) Prior heat stimulus sensitizes larvae to noxious mechanical inputs. (I) Control larvae (w^1118^) received a 10 s thermal stimulus on a peltier plate, and following 30 s of recovery received a single stimulus with a 50 mN von Frey filament. (J) Roll probability of control larvae following mechanical stimulus. *P < 0.05 Fisher’s exact test with BH correction. Sample numbers are indicated in each plot.

Next, we characterized the duration of this heat-evoked nociceptive sensitization. We presented larvae with two successive 41.5°C stimuli spaced by recovery periods ranging from 2 s to 5 min and assayed for reduced rolling latency to the second stimulus (Fig. 1D). We found that a prior noxious stimulus yielded sensitization that was apparent within 2 s of the initial stimulus (latency: 8.79 ± 3.36 s for first stimulus, 3.40 ± 2.12 s for second stimulus, n = 38; Fig. 1E); we could not reliably test shorter recovery periods because rolling responses often required >1 s for completion. We also observed sensitization with longer recovery periods (Fig. 1E), with significant heat-induced sensitization persisting for more than one minute (τ = 94.3 s, Fig. 1F). These data indicate that *Drosophila* larvae exhibit heat-gated nociceptive sensitization, which lasts for at least one minute and occurs primarily in the 30°C–45°C range.

*Drosophila* larvae exhibit mechanically-induced nociceptive sensitization that is triggered by epidermal responses to mechanical inputs^23^. To determine whether heat-induced nociceptive sensitization likewise reflects contributions of prior epidermis stimulation, we selectively stimulated epidermal cells or nociceptors using the red-shifted light-activated cation channel CsChrimson^33^ and assayed for sensitization to a subsequent noxious thermal stimulus (Fig. 1G). We found that optogenetic epidermal stimulation significantly sensitized larvae to a subsequent thermal stimulus, and that this sensitization was dependent on the obligate CsChrimson co-factor ATR (Fig. 1H). In contrast, we observed no significant effect of prior optogenetic stimulation on heat-induced rolling responses in effector-only controls or larvae expressing CsChrimson selectively in nociceptors. These data indicate that epidermal cells, not the nociceptor or its downstream circuitry, can trigger heat-induced nociceptive sensitization.

Prior studies demonstrated that optogenetic epidermal stimulation sensitizes larvae to noxious mechanical inputs^23^, therefore we examined whether thermal stimuli had a similar effect. Indeed, we found that prior exposure to heat inputs sensitizes larvae to subsequent noxious mechanical inputs. As was the case with temperature-evoked sensitization to noxious heat (Fig 1B, 1C), we observed a significant enhancement of nocifensive responses to mechanical inputs following prior exposure to 35°C and 40°C thermal stimuli (Fig. 1J).

### *Drosophila* larval epidermal cells are intrinsically thermosensitive

Our behavior studies demonstrate that epidermal cells drive heat-evoked sensitization to noxious temperature inputs, therefore we investigated whether larval epidermal cells are intrinsically responsive to environmental temperature changes. To this end, we developed a perfusion system that could deliver thermal ramps from ∼20°–45°C to larval specimens (Fig. 2A) and monitored responses to temperature change in larvae selectively expressing the Ca^2+^ indicator GCaMP6s^34^ in epidermal cells. *Drosophila* epidermal cells are sensitive to mechanical stimuli^23^, however perfusion of room temperature saline in our experimental paradigm did not induce significant epidermal GCaMP responses (Fig. 2B). We assayed epidermal responses to application (heat-on) and removal (heat-off) of noxious heat, targeting a temperature maximum of ∼42°C which yielded nociceptive sensitization in our behavior studies. In our thermal paradigm we observed no significant warming-induced increase in epidermal GCaMP6s signal (Fig. 2C). Instead, warming stimuli attenuated the GCaMP6s signal, consistent with prior reports of heat-induced reduction of GCaMP6s fluorescence^35^ and suggesting that epidermal cells do not exhibit robust responses to application of noxious heat. In contrast, epidermal cells exhibited a rapid and robust heat-off response; cooling from ∼42°C to 20°C induced a significant increase in GCaMP6s fluorescence that was apparent across the body wall epidermis (GCaMP6s ΔF/F_0_: 0.49 ± 0.47% for 20°C fluid flow; 82.34 ± 8.84% for heat off, n = 20 each) (Fig. 2C, 2D). Heat-evoked nociceptive sensitization persisted for >1 min after removal of heat stimulus (Fig. 1E, 1F); likewise, we found that epidermal heat-off responses persisted for > 2 min after the temperature returned to baseline and showed significant attenuation between 60 and 120 s (Fig. 2E; τ = 140.2 s, Supplementary Fig. 2).

**Figure 2.**
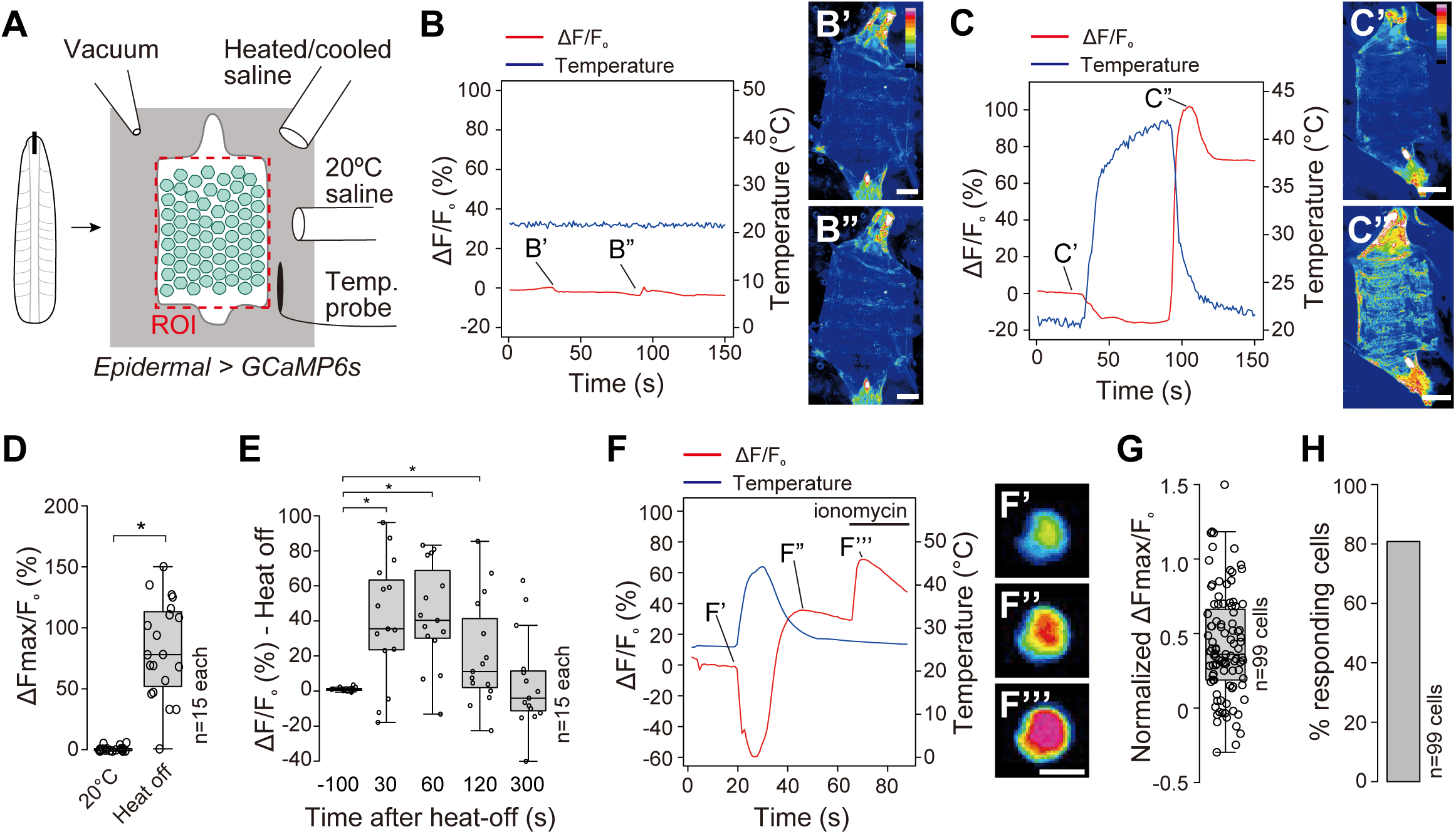
Epidermal cells are intrinsically thermosensitive. (A) Schematic of perfusion setup. Fillet preparations of larvae expressing the calcium reporter GCaMP6s in epidermal cells were pinned on sylgard pads within an imaging chamber, a temperature probe was placed adjacent to the specimen, and heated/cooled saline was perfused into the chamber to achieve temperature ramps. (B) Fluid flow alone does not trigger epidermal calcium responses. The plot depicts the GCaMP6s response (`Δ`F/F0, red line) and the solution temperature within the imaging chamber (blue line) for a representative specimen during perfusion with 20°C saline. Images depict epidermal GCaMP6s fluorescence at the indicated timepoints. (C) The larval epidermis responds to heat-off stimuli. The plots depict GCaMP6s responses of a representative larva to heating and heat-off stimuli. Images depict epidermal GCaMP6s fluorescence before application of heat stimulus (C’) and after removal of heat stimulus (C”). (D) Epidermal cells exhibit a robust heat-off response. Box plot depicts the maximum change in GCaMP6s fluorescence induced by perfusion of 20°C saline or by heat-off stimulus. *P < 0.05, Wilcoxon rank sum test. (E) Box plot depicts epidermal GCaMP6s fluorescence at the indicated timepoints for larval fillets that receive heat-off stimuli. n = 15 independent larvae each. *P < 0.05, Wilcoxon rank sum test with BH correction. (F-H) Heat-off responses of individual dissociated epidermal cells. (F) Traces depict temperature ramp and response of a representative individual epidermal cell to heating and heat-off stimuli. Ionomycin was added at the end of the temperature ramp to assess cell viability. Images depict GCaMP6s fluorescence at the indicated timepoints. Plots depict the amplitude of the GCaMP6s response (G) and proportion of cells that exhibited heat off responses (H). Genotype: *R38F11-GAL4, UAS-GCaMP6s*.

To determine whether these responses reflect an intrinsic responsiveness of epidermal cells to heat-off stimuli, we monitored temperature-evoked responses of isolated epidermal cells. Briefly, we dissociated the body wall of larvae expressing GCaMP6s in epidermal cells, immobilized the single cell suspension on poly-L-coated coverslips, and subjected cells to the thermal ramp stimulus that evoked heat-off responses in larvae. As with the semi-intact preparations, dissociated epidermal cells exhibited no significant response to fluid flow alone (Fig. 2F, prior to thermal stimulus).

Furthermore, we found that most epidermal cells exhibited heat-off responses, but not responses to warming or to noxious heat (Fig. 2F-2H). Taken together, these results indicated that the larval epidermis is intrinsically responsive to environmental temperature change, particularly to cooling after exposure to noxious heat.

### Epidermal heat off response depends on warming to a threshold temperature

The heat-off response we observed could reflect epidermal responses to cooling from a defined threshold or a more general response to temperature decrements. In the former scenario temperature ramps that did not reach the threshold temperature should not evoke responses, whereas the latter scenario predicts that epidermal cells would respond to relative temperature change regardless of the reference. To discriminate between these possibilities, we assayed for heat-off responses to thermal ramps with different baseline and maximum temperatures in semi-intact larval fillet preparations.

First, we fixed the maximum temperature of our thermal ramps at ∼40°C and assayed effects of reducing the magnitude of the temperature change (ΔT) by altering the baseline (Fig. 3A). Compared to our initial paradigm in which we started from a baseline of 20°C and warmed to ∼40°C to generate a ΔT of 20°C before cooling, we found that increasing the baseline temperature to 30°C while targeting the same maximum temperature still generated a robust heat-off response; the 50% reduction in ΔT was accompanied by only a ∼20% decrease of GCaMP6s ΔF/F_0_ signal (84.30 ± 6.00% vs 56.23 ± 5.15%, n=14, 15 each) (Fig. 3A, 3B). Next, we assayed effects of reducing both the baseline and maximum temperatures by 10°C to maintain a ΔT of 20°C. In this paradigm, epidermal cells exhibited negligible heat-off responses (ΔF/F_0_ = 1.92 ± 0.78%, n = 15 specimens) (Fig. 3A, 3B), suggesting that epidermal cells are responsive to cooling from a temperature threshold rather than the magnitude of temperature change.

**Figure 3.**
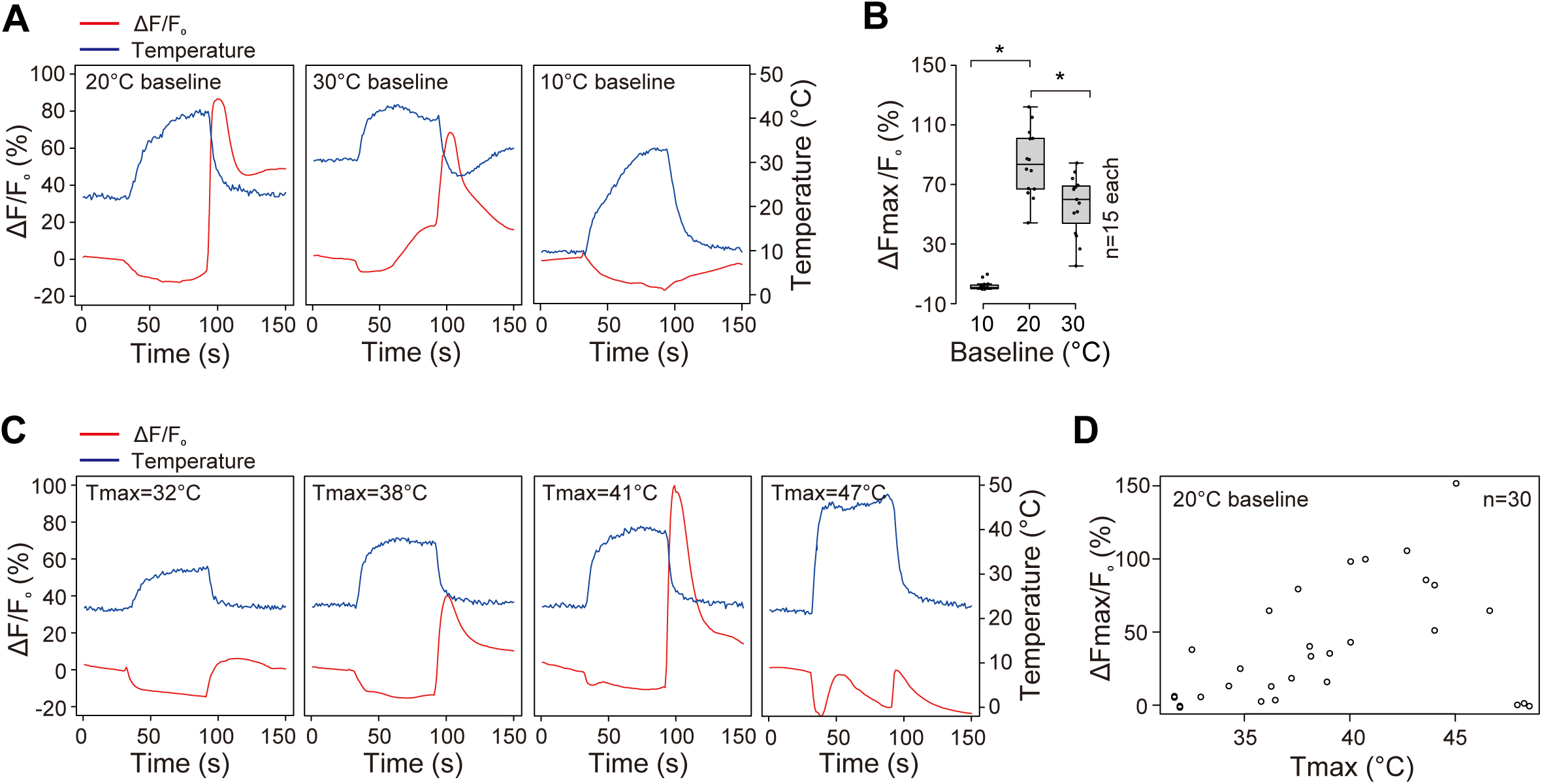
Epidermal cells exhibit a threshold-dependent heat off response. (A-B) Epidermal heat-off response is not a general response to cooling. (A) Representative traces depicting epidermal calcium responses to three different heat-off paradigms: left, 20°C baseline, 42°C max; middle, 30°C baseline, 42°C max; right, 10°C baseline, 32°C max. (B) Quantification of epidermal GCaMP6s responses to the heat-off paradigms depicted in (A). *P < 0.05, Kruskal-Wallis test followed by Wilcoxon rank sum test with BH correction. n = 15 larvae for each genotype. (C-D) Effects of temperature maxima on epidermal heat-off responses. (C) Traces depict epidermal GCaMP responses to heat-off responses to thermal ramps with a baseline of 22°C and maxima of 32°C, 38°C, 41°C, and 47°C. (D) Scatterplot depicts epidermal GCaMP6 responses to thermal ramps with a baseline of 22°C and a range of temperature maxima. Each circle represents an independent larval specimen, n = 30 larvae total. Genotype: *R38F11-GAL4, UAS-GCaMP6s*.

To identify the temperature threshold for the epidermal heat-off response we fixed the baseline temperature at 20°C and assayed epidermal responses to thermal ramps that targeted maximum temperatures ranging from 30°C–50°C (Fig. 3C, 3D). These studies revealed three general temperature-dependent aspects of the heat-off response. First, at temperature maxima below 35°C epidermal cells rarely exhibited heat-off responses (Fig. 3C, 3D). Second, at temperature maximum values between 35°C and 45°C, the magnitude of the heat-off response increased with increasing temperature. Third, at temperature maxima above 45°C the heat-off response was strongly attenuated, suggesting that the physiological response range of the larval heat off response lies between 35°C and 45°C. Altogether, these data indicate that the epidermal heat-off response is primarily dependent on warming to an absolute temperature; the relative change in temperature has a smaller contribution to the overall response.

Finally, to examine whether the heat-off responses reflect a general response to cooling in epidermal cells, we assayed epidermal calcium responses to cooling from room temperature rather than cooling after exposure to noxious heat. We found that epidermal cells exhibited a reproducible response to cooling in this paradigm (Supplementary Fig. 3). However, several observations suggest that this epidermal cooling response is distinct from the heat-off response. First, epidermal heat-off responses exhibited an exponential rise in GCaMP6s signal, whereas cooling responses were more gradual. Second, recovery from cooling exhibited more rapid linear decay kinetics (τ = 140.2 s for heat-off, 25.2 s for cooling). Third, the amplitude of cooling responses was dependent on the temperature minimum (Supplementary Fig. 3) whereas the heat-off response was dependent on the temperature maximum (Fig. 3D). Hence, epidermal cells exhibit distinct responses to heat-off and cooling stimuli.

### The heat-sensitive protein Stim is required for the epidermal heat-off response

Next, we sought to identify the molecular mechanisms responsible for the epidermal heat-off response. The increase in cytosolic calcium that we observed during the heat-off response could reflect entry of extracellular calcium into epidermal cells and/or release from intracellular stores. To distinguish between these possibilities, we monitored heat-off responses of larval skin preparations using Ca^2+^-free saline (0 mM Ca^2+^, 1.5 mM EGTA). Under these conditions, the epidermal GCaMP6s responses were significantly attenuated (ΔF/F_0_: 75.99 ± 5.88% for 1.5 mM Ca^2+^, 2.94 ± 0.62% for 0 mM Ca^2+^; p = 1.28 x 10^-8^, n=15 each) (Fig. 4A, 4B). Hence, the heat-off response in epidermal cells requires external calcium.

**Figure 4.**
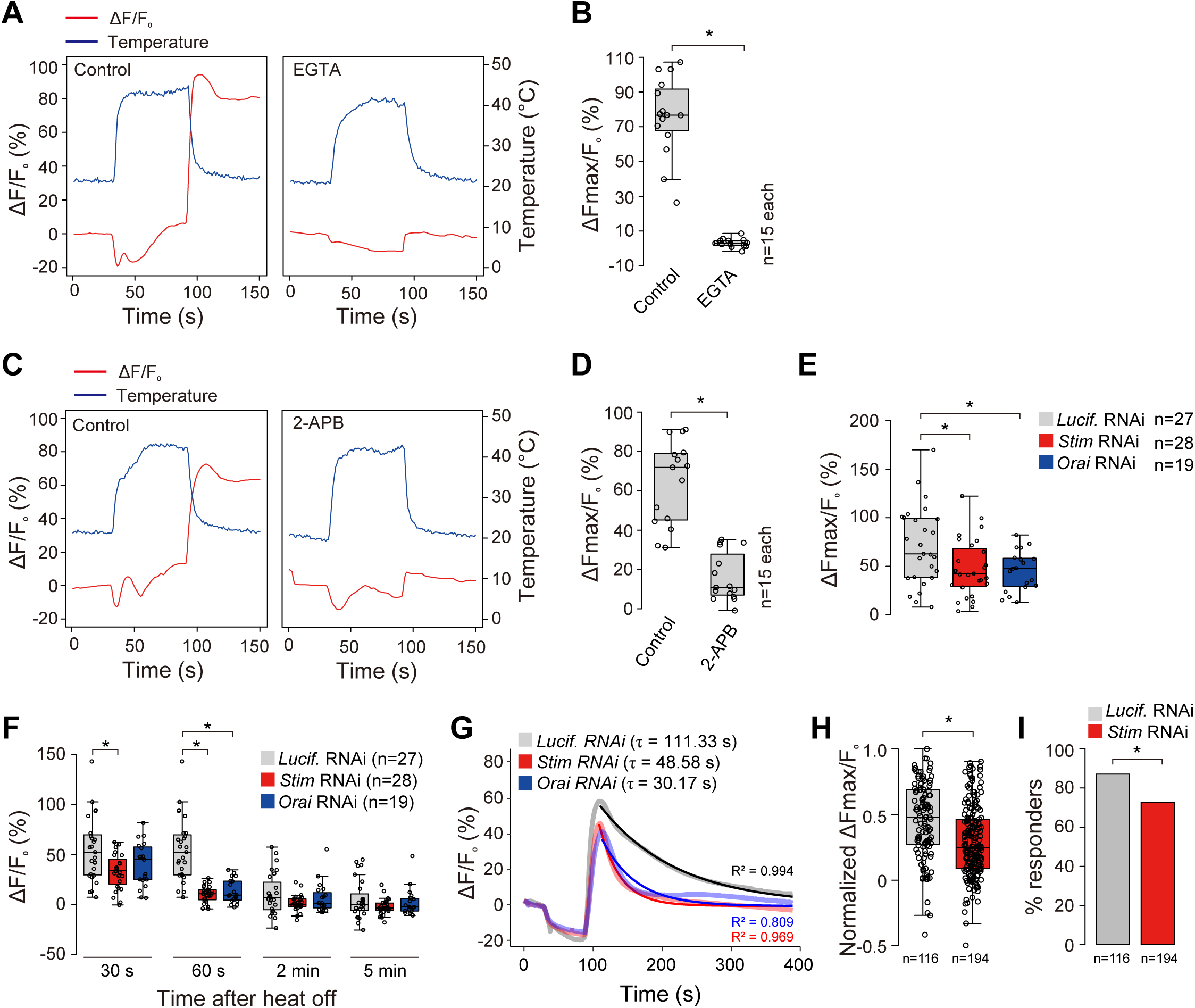
Epidermal heat-off responses require Stim and the CRAC channel Orai. (A-B) Epidermal heat-off responses require extracellular calcium. (A) Representative traces and (B) quantification of epidermal GCaMP6s responses to heat-off stimulus delivered in the absence or presence of 1.5 mM EGTA. *P < 0.05, Wilcoxon rank sum test. (C-D) The channel blocker 2-APB inhibits the epidermal heat-off response. (C) Representative traces and (D) quantification of epidermal GCaMP6s responses to heat-off stimulus delivered in the absence or presence of 100 mM 2-APB, a concentration that blocks Orai. GCaMP6s responses are significantly attenuated but not completely eliminated. *P < 0.05, Wilcoxon rank sum test. (E-G) Stim and Orai are required for heat-off response in *Drosophila* epidermal cells. (E) Maximum amplitude and (F) time course of epidermal GCaMP6s responses to heat-off stimulus are shown for larvae expressing *Luciferase* (control), *Stim*, or *Orai* transgenes in epidermal cells. *P < 0.05, Wilcoxon rank sum test with BH correction. (G) RNAi of *Stim* or *Orai* alters the kinetics of heat-off responses in epidermal cells. Traces indicate mean response profiles (light shading) and exponential curves (dark shading) fit to the decay profile of larvae of the indicated genotypes. (H-I) Stim is required for heat-off responses in dissociated epidermal cells. Plots depict epidermal GCaMP6s response amplitude (H) and fraction of cells that responded (I) to heat off stimuli for cells of the indicated genotypes. *P < 0.05, Wilcoxon rank sum test for H and Fisher’s exact test for I. The number of cells is indicated for each genotype. Genotypes: (A-D) *R38F11-GAL4, UAS-GCaMP6s*, (E-I) control: *R38F11-GAL4, UAS-Luciferase-RNAi, R38F11-LexA, LexAOP-GCaMP6s / +*; Orai RNAi: *R38F11-GAL4, UAS-Orai-RNAi, R38F11-LexA, LexAOP-GCaMP6s / +*; Stim RNAi: *R38F11-GAL4, UAS-Stim-RNAi, R38F11-LexA, LexAOP-GCaMP6s / +*.

Calcium release-activated calcium channels (CRAC), which are comprised of the Ca^2+^-selective cation channel Orai and its activator the transmembrane Ca^2+^ binding protein Stim, are required for heat-off responses in human and mouse keratinocytes^30^. Additionally, Stim is activated by temperature paradigms comparable to those that evoke heat-off responses in *Drosophila* epidermal cells: warming beyond 37°C followed by cooling^36^. We therefore investigated whether CRAC channels are required for the heat-off response in *Drosophila* epidermal cells. First, we monitored epidermal Ca^2+^ responses to thermal stimuli in the presence of the chemical inhibitor 2-aminoethoxydiphenyl borate (2-APB) at a concentration that blocks Orai as well as several TRP channels^37^. We found that 2-APB application significantly attenuated the epidermal heat-off response compared with vehicle-only controls (GCaMP6s ΔF/F_0_: 64.04 ± 5.49% for vehicle controls, 16.05 ± 3.20% for 100 mM 2-APB; p = 8.64 x 10^-7^, n=15 each) (Fig. 4C, 4D). To directly assay the requirement for *Stim* and *Orai* in the epidermal heat-off response we used RNAi to inactivate each gene in epidermal cells and monitored GCaMP6s responses to thermal stimuli (Fig. 4E). Indeed, we found that epidermal inactivation of either *Stim* or *Orai* by RNAi attenuated the amplitude and kinetics of the heat-off response, promoting a more rapid return to baseline calcium levels compared to controls (Fig. 4E, 4F). Likewise, we found that epidermal *Stim* RNAi significantly attenuated the proportion of dissociated epidermal cells that responded to heat-off stimuli (87.1% for control RNAi vs 72.7% for *Stim* RNAi) as well as the amplitude of the response (0.46 ± 0.027 for control RNAi vs 0.29 ± 0.02 for *Stim* RNAi; p < 0.05; n=116, 194 respectively) (Fig. 4G, 4H). Altogether, these results indicate that Stim and Orai are required cell-autonomously in epidermal cells for heat-off responses, which likely involves other channel(s) as well.

### The epidermal heat off response is essential for heat gated sensitization

The thresholds required to trigger heat-evoked nociceptive sensitization (Fig. 1) and epidermal heat-off responses (Fig. 3) were remarkably similar, therefore we investigated contributions of epidermal CRAC channels to epidermal heat-induced nociceptive sensitization (Fig. 5A). Like wild-type controls, larvae expressing a control RNAi transgene in epidermal cells exhibited a significant reduction in rolling latency to the second of two successive noxious thermal stimuli (Fig. 5B). In contrast, epidermal expression of *Stim* or *Orai* RNAi eliminated the heat-induced nociceptive sensitization without affecting the latency of the first response (*Stim* RNAi roll latency: 7.16 ± 0.91 s, 1^st^ stimulus; 6.57 ± 0.83 s, 2^nd^ stimulus; n = 34 larvae) (Fig. 5B). These data indicate that epidermal CRAC channels are essential for heat-gated nociceptive sensitization.

**Figure. 5.**
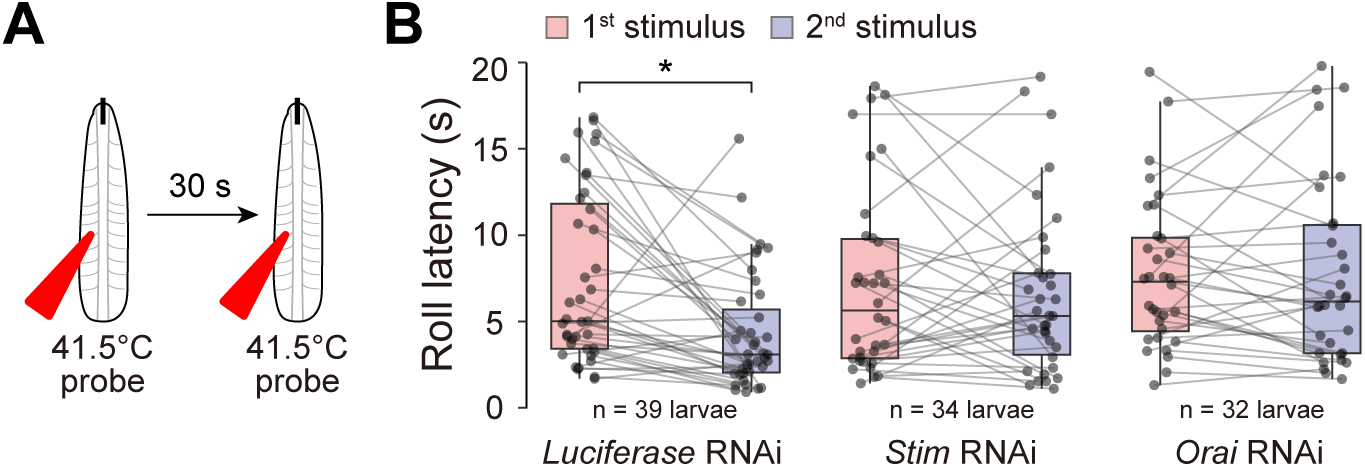
CRAC channels are required for epidermal heat-evoked nociceptive sensitization. (A) Schematic of experimental paradigm. Control larvae (*w^1118^*) received two successive 41.5°C stimuli separated by a 30 s recovery period. (B) Plots depict the roll latency to each of the 41.5°C stimuli for larvae expressing Luciferase (control), Stim, and Orai RNAi selectively in epidermal cells. *P < 0.05, Wilcoxon rank sum test with BH correction. Genotypes: control: *R38F11-GAL4, UAS-Luciferase-RNAi / +*; *Orai* RNAi: *R38F11-GAL4, UAS-Orai-RNAi / +*; *Stim* RNAi: *R38F11-GAL4, UAS-Stim-RNAi / +*.

## Discussion

In this study, we defined a role for *Drosophila* epidermal cells in escape responses to noxious heat. Prior heat exposure sensitizes larvae to noxious heat, and we find that epidermal but not nociceptor activation yields this form of noxious heat sensitization. Our work defines several features of this sensitization and extends prior work defining epidermal thermosensory responses. First, the sensitization is short-lived; prior heat exposure primes larvae to future inputs that occur within seconds to minutes, potentiating escape from a harmful environment without providing indefinite sensitization to noxious inputs that would be maladaptive. Second, the sensitization is increasingly triggered by warm temperatures outside of the preferred range. Hence, as temperatures of prior stimuli approach noxious levels, so too does the state of vigilance they induce. Third, epidermal thermosensory responses that underlie the sensitization are heat-off responses, suggestive of a mechanism by which epidermal cells reinforce productive escape behavior. Prior studies demonstrated that CRAC channels mediate heat-off responses in epidermal cells and that epidermal CRAC channel inactivation affected thermal preference^30^; our studies demonstrate that epidermal CRAC channels are required for epidermal heat-induced sensitization of nociceptors. Fourth, prior heat stimulus yields polymodal sensitization: responses to noxious thermal as well as mechanosensory inputs, suggesting that epidermal heat-evoked signals broadly sensitize nociceptors to future stimuli. The heat-evoked sensitization we describe here is remarkably similar to mechanically-evoked nociceptive sensitization seen in *Drosophila* larvae, which likewise emerges on a timescale of seconds and yields reversible, mechanical hypersensitivity to protect larvae from further insult^23^. Heat-induced sensitization to noxious inputs has been reported in *Manduca* larvae as well^38^, but epidermal contributions have not yet been explored. These forms of sensitization are distinct from pathological forms of nociceptive sensitization that emerge on a timescale of hours, and are long-lasting^13,16,39^.

Our studies suggest that epidermal cells exhibit robust responses to heat-off stimuli as well as responses to cooling stimuli but negligible responses to heat-on stimuli. This result is somewhat surprising given the prior findings that mouse keratinocytes exhibited both heat-on and heat-off responses^29,30^ and that *Drosophila* epidermal cells express several temperature-gated ion channel genes including TRPA (*painless* and *pyrexia*) and TRPM (*Trpm* and *Trpml)* channels^23^. What is the basis of this selectivity? Stim/Orai are required for both the heat-off response of epidermal cells and the resulting nociceptive sensitization. Stim is a transmembrane protein that functions as a sensor for ER calcium concentration and an activator of the calcium-selective cation channel Orai. Stim also functions as a temperature sensor with properties similar to the response properties we observed in epidermal cells: warming beyond a threshold of ∼39°C transitions Stim to the active state and drives clustering ER-PM contact sites, but subsequent cooling (“heat-off”) is required for Orai opening^36,40^ In addition to its function as a thermosensor, Stim is required for mechanosensory responses of both *Drosophila* epidermal cells and human keratinocytes^23^. Intriguingly, STIM1 can also be activated by oxidative stress, with S-glutathionylation promoting unfolding of the EF-SAM calcium binding domain^41^. Hence, Stim may be a nexus for sensory processing in the skin.

What are the outputs of the Stim/Orai-mediated rise in epidermal calcium that induce nociceptor sensitization? Mechanically-induced nociceptor sensitization triggered by epidermal mechanosensory responses likely requires vesicular release, as the sensitization is significantly attenuated by acute blockade of exocytosis using a dominant-negative temperature sensitive allele of *shibire*, the gene encoding *Drosophila* dynamin^23^. The temperature thresholds required to trigger heat-evoked sensitization precluded a similar analysis of acute requirements for epidermal dynamin function in the process. However, given the similarity in timescales of mechanically evoked and heat-evoked nociceptor sensitization, the shared requirement for epidermal Stim and Orai, and the observation that epidermal stimulation potentiates nociceptors to multimodal inputs, it seems likely that both forms of potentiation rely on common signaling output.

In mammals, keratinocytes release a variety of substances that potentiate nociceptor function including ATP, inflammatory mediators, and neuropeptides. Among these, genetic and pharmacological evidence support a role for ATP signaling in mechanically-and possibly heat-evoked nociceptor sensitization^21,22,29^. Although *Drosophila* lacks classical purinergic receptors, nociceptors express Adenosine receptor (AdoR), and gliotransmission appears to sensitize nociceptors via AdoR signaling following injury^42^. AdoR is expressed in nociceptors but no other SSNs, hence ATP/adenosine release would be one mechanism to selectively couple epidermal cells to nociceptors. However, AdoR is coupled to adenylate cyclase, which is unlikely to work on the timescales we observe. In the mammalian somatosensory system, inflammatory mediators and neuropeptides promote nociceptor hypersensitivity via TRPA1 and/or TRPV1 channel activation^43,44^, hence it will be intriguing to determine whether epidermal nociceptive sensitization is mediated by TRP channels in nociceptors.

In addition to the heat-off responses, *Drosophila* epidermal cells respond to cooling from comfortable to cold temperatures. These cooling responses differed from the Stim/Orai-dependent heat off responses in their kinetics and temperature dependance, hence they likely involve different molecular mediators. Brief exposure to chilling in a variety of insects triggers an adaptive response referred to as rapid cold hardening (RCH), which is mediated by a rise in intracellular calcium in peripheral tissues^45^. RCH likely occurs as a result of cold-inhibition of transporters that maintain calcium homeostasis^46,47^, and is induced most effectively by temperatures between 0°C and 10°C, comparable to the threshold we observed for cooling-dependent calcium influx in epidermal cells^48,49^. Although the timescales differ somewhat, it seems plausible that the cooling responses we observed are mechanistically linked to RCH.

Alternatively, epidermal cells may express cool sensors that mediate these responses, and indeed mRNA for Trpm and Trpml, which contribute to larval responses to noxious cold are expressed in epidermal cells^6,50^. It will therefore be intriguing to determine whether epidermal calcium responses to cooling contribute to nociceptor sensitization observed during cold acclimation^51^.

## Methods

| Key Resources Table |  |  |  |  |
| --- | --- | --- | --- | --- |
| Reagent type (species) or resource | Designation | Source or reference | Identifiers | Additional information |
| gene (*D. melanogaster*) | Stim | GenBank | FLYB:FBgn0045073 |  |
| gene (*D. melanogaster*) | Orai | GenBank | FLYB:FBgn0041585 |  |
| genetic reagent (*D. melanogaster*) | *GMR38F11-GAL4* | Bloomington Drosophila Stock Center | BDSC_50014 | GAL4 driver (epidermis) |
| genetic reagent (*D. melanogaster*) | *GMR38F11-LexA* | This study | This study | LEXA driver (epidermis) |
| genetic reagent (*D. melanogaster*) | *ppk-GAL4* | Bloomington Drosophila Stock Center | BDSC_32079 | GAL4 driver (C4da neurons) |
| genetic reagent (*D. melanogaster*) | *ppk-LexA* | Maintained in the Parrish Lab | Flybase_FBtp0125814 | LEXA driver (C4da neurons) |
| genetic reagent (*D. melanogaster*) | *UAS-GCaMP6s* | Bloomington Drosophila Stock Center | BDSC_42746 | Calcium indicator |
| genetic reagent (*D. melanogaster*) | *AOP-GCaMP6s* | Bloomington Drosophila Stock Center | BDSC_44273 | Calcium indicator |
| genetic reagent (*D. melanogaster*) | *UAS-CsChrimson* | Bloomington Drosophila Stock Center | BDSC_55136 | Inducible Channel |
| genetic reagent (*D. melanogaster*) | *UAS-Luciferase RNAi* | Bloomington Drosophila Stock Center | BDSC_31603 | RNAi transgene |
| genetic reagent (*D. melanogaster*) | *UAS-orai-RNAi* | Bloomington Drosophila Stock Center | BDSC_53333 | RNAi transgene |
| genetic reagent (*D. melanogaster*) | *UAS-stim-RNAi* | Bloomington Drosophila Stock Center | BDSC_41759 | RNAi transgene |
| recombinant protein | Collagenase type I | Thermo Fisher | Cat. #: 17-100-017 |  |
| chemical compound | All-trans retinal | Millipore Sigma | Cat. #: R2500 |  |
| chemical compound | Lanthanum chloride | Millipore Sigma | Cat. #: 211605 |  |
| chemical compound | ionomycin | Sigma-Aldrich | Cat. #: I0634 |  |
| chemical compound | Poly-L-lysine | Millipore Sigma | Cat. #: P1524 |  |
| software, algorithm | ImageJ | ImageJ | RRID:SCR_003070 |  |
| software, algorithm | R | R Foundation | RRID:SCR_001905 |  |
| software, algorithm | Metafluor | Molecular Devices | RRID:SCR_014294 |  |
| software, algorithm | NIS-Elements | Nikon | RRID:SCR_014329 |  |
| software, algorithm | pClamp | Molecular Devices | RRID:SCR_011323 |  |

### Drosophila strains

Flies were maintained on standard cornmeal-molasses-agar media and reared at 25°C under 12 h alternating light-dark cycles. Third-instar larvae (96-120 AEL) larvae were used in all experiments. All alleles except for *R38F11-LexA* were obtained from the Bloomington Drosophila Stock Center. To generate the *R38F11-LexA* driver line (*R38F11-LexA-GAD*), we used LR clonase (Thermo Fisher) to shuttle the *GMR38F11* enhancer from a Gateway cloning vector (Rubin lab) to pBPnlsLexA::GADflUw^52^. The resulting expression vector (pBP-R38F11-nlsLexA::GADflUw) was integrated at the VK00027 attP docking site (BDSC_9744) by BestGene.

### Calcium imaging of fillet preparations

Third-instar larvae were dissected along the ventral midline and pinned on sylgard (Dow Corning) dishes with the internal surface facing up. All internal organs, including the central nervous system, were removed. Larvae were bathed in Fly Saline Solution (Supplementary Table 1). Solution was heated using a CL-100 Temperature Controller (Warner Instruments) and delivered to imaging chambers through an SC-20 inline Peltier heater (Warner Instruments) using a pinch valve perfusion system (Automate Scientific). Temperature was captured using a BAT-12R thermocouple (Physitemp) placed next to the specimen in the imaging chamber, and temperature readings were digitized using a DAQ device (USB-6529, National Instruments). Data was acquired by Metafluor (Molecular Device) and analyzed by Image J^53^. For curve fitting, sample-averaged fluorescence traces were fitted with a single exponential decay function using R to extract a representative time constant (τ) and assess response kinetics

### Calcium imaging of dissociated cells

Twelve larval fillets were dissociated in 500 µL of PBS with 200 U/mL collagenase type I (Thermo Fisher), with mixing at 1000 RPM at 37°C for 15 min, with trituration every 5 min. Undigested fillets were removed and the remaining suspension was spun at 500 x g for 3 min, followed by aspiration of the supernatant down to a 10 µL cell suspension. Cells were resuspended in 50 µL of a 1:1 solution of fly saline (Supplementary Table 1) and Schneider’s insect media (Thermo Fisher) solution and plated onto poly-D-lysine (1 mg/ml, Merck P6407) coated 12-mm coverslips, with 10 µL cell solution per coverslip. TE300 microscope (Nikon) equipped with CoolSNAP DYNO (Photometrics) was used to perfuse a heated solution. A heated bath solution was perfused, followed by 5 μM ionomycin (Millipore Sigma). Images and temperature were captured by NISElement AR (Nikon) and AxoScope (Molecular Devices), respectively. All data were displayed as a normalized ratio relative to ionomycin.

### Behavior assays

Third instar larvae were isolated from their food, washed in distilled water, and placed on a scored 35 mm petri dish with a thin film of water such that larvae stayed moist but did not float. For thermonociception assays, larvae were stimulated dorsally between segments A4 and A7 with a thermal probe (ProDev Engineering, Missouri City, TX) set to 41.5°C, a temperature that yielded nocifensive rolling responses in >75% of larvae with 8-10 s of stimulus on average^32^, allowing for identification of treatments that sensitized (reduced latency) or desensitized (increased latency) larvae to noxious heat.

The probe was held in constant contact with the larval skin for 20 s or until a nocifensive roll was completed, larvae were allowed to recover during free locomotion for the indicated interval, and a second 20 s thermal stimulus was applied. Behavior Responses were analyzed post-hoc blind to genotype and were plotted according to roll probability and roll latency.

For assays testing effects of varying the temperature of prior thermal stimuli on thermal nociception, larvae were individually transferred to a pre-warmed Peltier plate containing a thin layer of water, incubated for 10 s at the indicated temperature, and transferred to the behavior arena. Following 10 s of recovery, larvae were stimulated with a 41.5°C thermal probe, as above, and latency to the first complete roll was recorded.

For assays involving thermal and mechanical stimuli, larvae were individually transferred to a pre-warmed Peltier plate containing a thin layer of water, incubated for 10 s at the indicated temperature, and transferred to the behavior arena with a paint brush. Following 10 s of recovery, larvae were mechanically stimulated between segments A4 and A7 with a calibrated Von Frey filament that delivered the indicated force upon buckling. Nocifensive rolling responses were scored during the 10 s following stimulus removal.

For assays involving optogenetic and mechanical stimuli, larvae were raised in constant dark at 25°C on food supplemented with 1 mM all-trans retinal, transferred individually to a 100 mM 2% agar plate, and habituated in the behavior arena for 30 s. Larvae were stimulated for 10 s with 300 μW/mm^2 broad-spectrum (500-700 nm) LED illumination (CoolLED PE-300, green), individually transferred to a 35 mM petri dish, as above, and assayed for thermal nociception responses 30 s after completion of optical stimulus.

### Statistical analysis

Statistical analyses were conducted using R software. Details of statistical tests including treatment groups, sample numbers, statistical tests, p-values and q-values are provided in the Supplementary Methods.

## Acknowledgements

Fly stocks obtained from the Bloomington Drosophila Stock Center (NIHP40OD018537) were used in this study. We thank Gerry Rubin for sharing vectors for fly transgenics, and the Parrish lab for critical input.

## Author Contributions

`●` Conceptualization: JY and JZP
`●` Methodology: JY, TM,TS and JZP
`●` Investigation: JY, AC, CY, FT and JZP
`●` Data curation: JY
`●` Formal Analysis: JY
`●` Visualization: JY
`●` Supervision: TS, KE and JZP
`●` Funding acquisition: KE and JZP
`●` Writing – original draft: JY and JZP
`●` Writing – review and editing: JY, FT, KE, TS, and JZP

---

**Supplementary Figure 1. Related to Figure 1.**
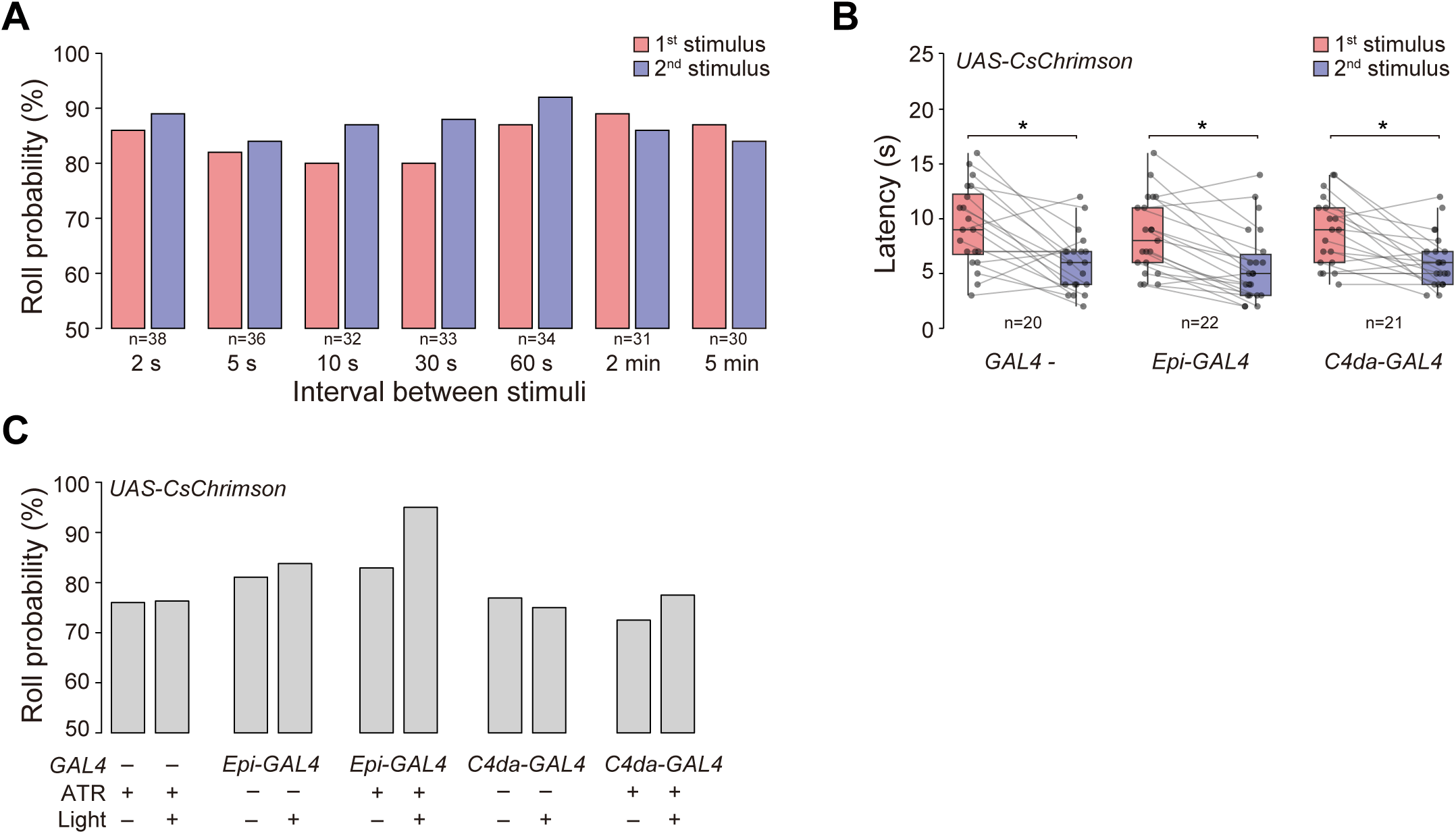
(A) Plot depicts roll probability of control larvae (*w^1118^*) in response to two successive 41.5°C stimuli separated by the indicated recovery periods. Roll latency values are shown in Fig. 1E. Genotype: *w^1118^*. (B) Experimental genotypes used in Figure 1G-1H exhibit heat-evoked nociceptive sensitization. Larvae received two successive 41.5°C stimuli separated by a 30 s recovery period and the roll latency values for each of the 41.5°C stimuli are shown in the plot. *P < 0.05, paired Wilcoxon rank sum test with BH correction. Genotypes: GAL4 − (*w^1118^; UAS-CsChrimson/+*), *Epi-GAL4* (*w^1118^; R38F11-GAL4, UAS-CsChrimson/+)*, and *C4da-GAL4* (*w^1118^; ppk-GAL4, UAS-CsChrimson/+*). (C) Companion to Fig. 1G. Plot depicts roll probability of effector controls (*w^1118^; UAS-CsChrimson/+*), larvae expressing CsChrimson in the epidermis (*w^1118^; R38F11-GAL4, UAS-CsChrimson/+)*, and larvae expressing CsChrimson in C4da neurons (*w^1118^; ppk-GAL4, UAS-CsChrimson/+*) in response to 41.5°C thermal stimulus. Larvae were raised on media containg (+ ATR) or lacking (-ATR) the obligate CsChrimson co-factor ATR, as indicated. Prior optogenetic epidermal stimulation did not significantly alter the roll probability in response to a 41.5°C thermal stimulus, Fisher’s exact test with BH correction.

**Supplementary Figure 2. Related to Figure 2.**
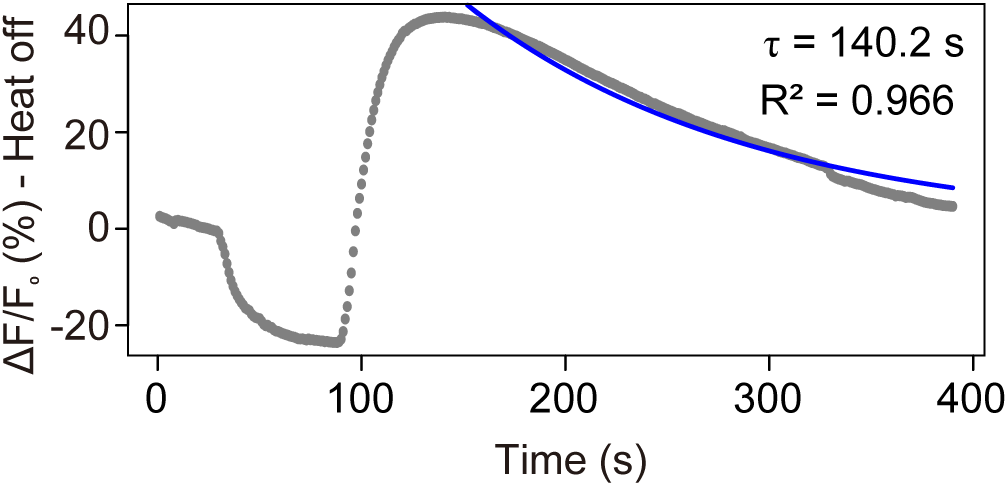
Time constant for heat-off response in larval epidermis. The decay time constant was calculated by fitting the data points from the peak response to the end of the experiment into an exponential curve f(x) = a*exp(b*x) using R software. The time constant for the decay (τ) = 140.2 s.

**Supplementary Figure 3. Related to Figure 3.**
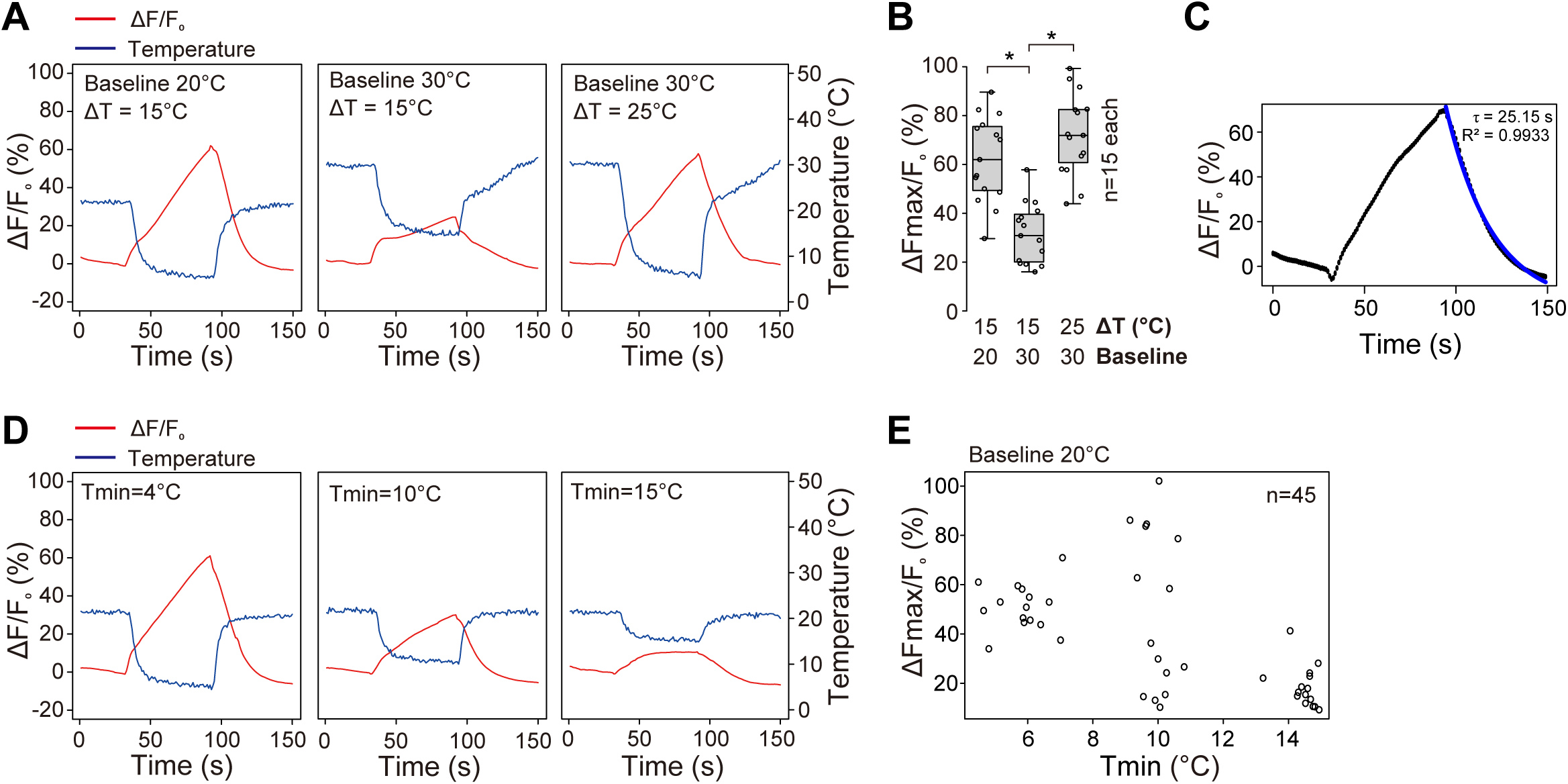
Epidermal cells exhibit a threshold-dependent cooling response. (A) Representative traces depicting epidermal calcium responses to three different cooling paradigms: left, 20°C baseline, 5°C min; middle, 30°C baseline, 15°C min; right, 30°C baseline, 5°C min. (B) Quantification of epidermal GCaMP6s responses to the heat-off paradigms depicted in (A). *P < 0.05, Kruskal-Wallis test followed by Wilcoxon rank sum test with BH correction. n = 15 larvae for each genotype. (C) Time constant for cooling response. The decay time constant was calculated by fitting the data points from the peak response to the end of the experiment into an exponential curve F(t) = Fmax⋅e^−t/τ^ using R software. The time constant for the decay (τ) = 25.15 s. (D-E) Effects of temperature minima on epidermal heat-off responses. (D) Traces depict epidermal GCaMP responses to heat-off responses to thermal ramps with a baseline of 20°C and minima of 4°C, 10°C, and 15°C. (E) Scatterplot depicts epidermal GCaMP6 responses to thermal ramps with a baseline of 20°C and a range of temperature minima. Each circle represents an independent larval specimen, n = 45 larvae total. Genotype: *R38F11-GAL4, UAS-GCaMP6s*.

**Supplementary Table 1**. Recipes for solutions used in imaging and physiology experiments.

