## Supplemental Methods for "Heat-off responses of epidermal cells sensitize *Drosophila* larvae to noxious inputs"

**Supplementary Table 1. Solutions used in calcium imaging studies.**

| **HL3.1, *Drosophila* saline** | | |
|---|---|---|
| **Compound** | **M.W. (g/mol)** | **1X Concentration (mM)** |
| NaCl | 58.44 | 120 |
| KCl | 74.55 | 5 |
| Proline | 115.13 | 5 |
| HEPES | 238.30 | 10 |
| Trehalose | 378.33 | 5 |
| Sucrose | 342.30 | 32.5 |
| $CaCl_2$ | 1M stock | 1.5 |
| $MgCl_2$ | 1M stock | 1 |
| pH (with NaOH) | 7.15 | |
| Osmolarity (mOsm/L) | 310 | |

| **Zero $Ca^{2+}$ / EGTA HL3.1** | | |
|---|---|---|
| **Compound** | **M.W. (g/mol)** | **1X Concentration (mM)** |
| NaCl | 58.44 | 120 |
| KCl | 74.55 | 5 |
| Proline | 115.13 | 5 |
| HEPES | 238.30 | 10 |
| Trehalose | 378.33 | 5 |
| Sucrose | 342.30 | 32.5 |
| EGTA | 380.35 | 1.5 |
| $MgCl_2$ | 1M stock | 1 |
| pH (with NaOH) | 7.15 | |
| Osmolarity (mOsm/L) | 310 | |

**Supplementary Methods: Details of Statistical Analysis**

**Fig. 1B**: Kruskal-Wallis test followed by Wilcoxon rank sum test with BH correction

| | p-value | q-value | Significance |
|---|---|---|---|
| Kruskal-Wallis test | 0.0002134 | | T |
| no heat vs 25 | 8.499027e-01 | 0.8499026568 | F |
| no heat vs 30 | 3.090032e-02 | 0.0386253958 | T |
| no heat vs 35 | 7.338206e-03 | 0.0183455161 | T |
| no heat vs 40 | 5.003549e-05 | 0.0002501775 | T |
| no heat vs 45 | 2.030552e-02 | 0.0338425319 | T |

**Fig. 1C**: Fisher's exact test with BH correction

| | p-value | q-value | Significance |
|---|---|---|---|
| no heat vs 25 | 1 | 1.000000 | F |
| no heat vs 30 | 0.6242 | 0.816125 | F |
| no heat vs 35 | 0.308 | 0.770000 | F |
| no heat vs 40 | 0.0226 | 0.113000 | F |
| no heat vs 45 | 0.6529 | 0.816125 | F |

**Fig. 1E**: paired Wilcoxon rank sum test with BH correction

| | p-value | q-value | Significance |
|---|---|---|---|
| 2 sec | 0.000000172 | 0.00000121 | T |
| 5 sec | 0.000101 | 0.000177 | T |
| 10 sec | 0.00000761 | 0.0000178 | T |
| 30 sec | 0.00000344 | 0.0000120 | T |
| 60 sec | 0.000214 | 0.000300 | T |
| 2 min | 0.0466 | 0.0544 | F |
| 5 min | 0.973 | 0.973 | F |

**Fig. 1H**: Wilcoxon rank sum test with BH correction

| UAS-CsCrimson+ | p-value | q-value | Significance |
|---|---|---|---|
| ATR+ light- vs opto stim | 0.7586743395 | 0.870133735 | F |
| 38F11-GAL4 ATR- light- vs opto stim | 0.8675698605 | 0.870133735 | F |
| 38F11-GAL4 ATR+ light- vs opto stim | 0.0000587867 | 0.000293934 | T |
| ppk-GAL4 ATR- light- vs opto stim | 0.6416227609 | 0.870133735 | F |
| ppk-GAL4 ATR+ light- vs opto stim | 0.8701337351 | 0.870133735 | F |

**Fig. 1J**: Fisher's exact test with BH correction

| | p-value | q-value | Significance |
|---|---|---|---|
| 25 vs 30 | 0.2437 | 0.24370000 | F |
| 25 vs 35 | 0.001299 | 0.00194850 | T |
| 25 vs 40 | 2.249e-05 | 0.00006747 | T |

**Fig. 2E**: Kruskal-Wallis test followed by Wilcoxon rank sum test with BH correction

| | p-value | q-value | Significance |
|---|---|---|---|
| Kruskal-Wallis test | 2.164e-10 | | T |

| | | | |
|---|---|---|---|
| No temp vs Heat off | 4.353e-10 | 4.35e-10 | T |
| No temp vs Cold | 1.451e-11 | 2.90e-11 | T |

**Fig. 2F**: Kruskal-Wallis test followed by Wilcoxon rank sum test with BH correction

| | p-value | q-value | Significance |
|---|---|---|---|
| Kruskal-Wallis test | 0.0002342 | | T |
| Before heat vs 30 sec | 0.004266 | 0.00711 | T |
| Before heat vs 60 sec | 0.000008819 | 0.000044095 | T |
| Before heat vs 2min sec | 0.0209 | 0.026125 | T |
| Before heat vs 5 min | 0.3892 | 0.3892 | F |

**Fig. 3B**: Kruskal-Wallis test followed by Wilcoxon rank sum test with BH correction

| | p-value | q-value | Significance |
|---|---|---|---|
| Kruskal-Wallis test | 8.148e-08 | | |
| RT vs 30℃ | 0.003153 | 3.153e-03 | T |
| RT vs 10℃ | 2.579e-08 | 5.158e-08 | T |

**Fig. 4B**: Wilcoxon rank sum test

| | p-value | q-value | Significance |
|---|---|---|---|
| Control vs EGTA | 1.289e-08 | | T |

**Fig. 4D**: Wilcoxon rank sum test

| | p-value | q-value | Significance |
|---|---|---|---|
| Control vs 2-APB | 8.639e-07 | | T |

**Fig4E**:Kruskal-Wallis test followed by Wilcoxon rank sum test with BH correction

| | p-value | q-value | Significance |
|---|---|---|---|
| Kruskal-Wallis test | 0.02938 | | T |
| Luci vs Stim | 0.02486 | 0.02486 | T |
| Luci vs Orai | 0.0205 | 0.02486 | T |

**Fig. 4F**: Kruskal-Wallis test followed by Wilcoxon rank sum test with BH correction

| | p-value | q-value | Significance |
|---|---|---|---|
| Kruskal-Wallis test | p-value< 2.2e-16 | | |
| 30sec Luci vs stim | 0.1434 | 0.2868000 | F |
| 30sec Luci vs Orai | 0.2711 | 0.3614667 | F |
| 60 sec Luci vs Stim | 0.0009504 | 0.0076032 | T |
| 60 sec Luci vs Orai | 0.002173 | 0.0086920 | T |
| 2 min Luci vs Stim | 0.1028 | 0.2741333 | F |

| | | | |
|---|---|---|---|
| 2min Luci vs Orai | 0.4329 | 0.4947429 | F |
| 5min Luci vs Stim | 0.2066 | 0.3305600 | F |
| 5min Luci vs Orai | 0.684 | 0.6840000 | F |

**Fig. 4G**: Wilcoxon rank sum test

| | p-value | q-value | Significance |
|---|---|---|---|
| Control vs Stim RNAi | 2.281e-07 | | T |

**Fig. 4H**: Fisher's exact test

| | p-value | q-value | Significance |
|---|---|---|---|
| Control vs Stim RNAi | 0.002895 | | T |

**Fig. 5B**: Wilcoxon rank sum test with BH correction

| | p-value | q-value | Significance |
|---|---|---|---|
| Control 1st vs control 2nd | 5.496e-05 | 0.00010992 | T |
| Stim RNAi 1st vs Stim RNAi 2nd | 0.6606 | 0.66060000 | F |

**Supp. Fig. 1A**: Kruskal-Wallis test and Wilcoxon rank sum test with BH correction

| | p-value | q-value | Significance |
|---|---|---|---|
| Kruskal-Wallis test Before | 0.68313 | | F |
| Kruskal-Wallis test After | 0.3684 | | F |
| Kruskal-Wallis test After-Before | 0.5724 | | F |
| UAS-CsChrimson Before vs After | 0.000948 | 0.0014220 | T |
| R38F11-GAL4, UAS-CsChrimson Before vs After | 0.0001926 | 0.0005778 | T |
| ppk-GAL4, UAS-CsChrimson Before vs After | 0.005081 | 0.0050810 | T |

**Supp. Fig. 1B**: Fisher's exact test with BH correction

| | p-value | q-value | Significance |
|---|---|---|---|
| 2s before vs after | 1.0000 | 1 | F |
| 5s before vs after | 1.0000 | 1 | F |
| 10s before vs after | 0.5458 | 1 | F |
| 30s before vs after | 0.5468 | 1 | F |
| 60s before vs after | 0.7115 | 1 | F |
| 2min before vs after | 1.0000 | 1 | F |
| 5min before vs after | 1.0000 | 1 | F |

**Supp. Fig. 1C**: Fisher's exact test with BH correction

| | p-value | q-value | Significance |
|---|---|---|---|
| No GAL4 light - vs | 1.0000 | 1 | F |

| | | | |
|---|---|---|---|
| light + | | | |
| 38F11GAL4 no ATR light - vs light + | 1.0000 | 1 | F |
| 38F11GAL4 ATR light - vs light + | 0.1549 | 0.7745 | F |
| ppkGAL4 noATR light - vs light + | 1.0000 | 1 | F |
| ppkGAL4 ATR light- vs light + | 0.7968 | 1 | F |

**Supp. Fig. 2**: Kruskal-Wallis test followed by Wilcoxon rank sum test with BH correction

| | p-value | q-value | Significance |
|---|---|---|---|
| Kruskal-Wallis test | 3.338e-06 | | T |
| RT vs 30 | 0.1485 | 0.1485000 | F |
| RT vs 30-15 | 1.555e-05 | 0.0000312 | T |
